# An indirect organogenesis-based citrus transformation system monitored using visible reporter markers

**DOI:** 10.64898/2026.09.05.749627

**Authors:** Sameena Ejaz Tanwir, Tao Jiang, Anandi Karn, Carlos D. Messina, Heqiang Huo

## Abstract

Genetic transformation of citrus is essential for functional genomics and trait improvement, yet remains limited by prolonged regeneration cycles, genotype-dependent responses, and inefficient screening of transformed tissues. Here, we established an *Agrobacterium*-mediated indirect organogenesis-based transformation system for Carrizo citrange and evaluated two visible reporter systems, the anthocyanin regulator *ROSEA1* and the betalain biosynthetic cassette *RUBY*, for monitoring transformed tissues throughout regeneration. An optimized regeneration protocol enabled visible callus initiation within 7 days following a 5-day pre-culture treatment and recovery of PCR-confirmed transgenic plantlets within 5 months. Both visible reporters were compatible with callus development and shoot regeneration throughout the indirect organogenesis workflow. During callus proliferation, visible pigmentation closely corresponded with GFP fluorescence, enabling rapid, non-destructive identification of transformed tissues. However, reporter performance diverged during shoot regeneration: *RUBY* maintained stable pigmentation throughout regeneration, whereas *ROSEA1*-associated pigmentation progressively declined despite continued GFP fluorescence. Consequently, *RUBY* exhibited a 4.7-fold higher pigmentation-based detection rate than *ROSEA1* at the shoot stage and showed closer agreement with GFP-based detection. Putative transgenic events were confirmed by PCR in independent lines of both constructs, together with GFP fluorescence in leaves and root tips. Together, these findings establish an efficient indirect organogenesis-based transformation platform for citrus and demonstrate that both *ROSEA1* and *RUBY* are effective visual reporters during callus proliferation, whereas *RUBY* provides more reliable visual identification during shoot regeneration and plant recovery.

## 1. Introduction

Citrus is one of the world’s most economically important fruit crops, with an annual production of approximately 161.8 million tons, and represents a major source of vitamin C, flavonoids, carotenoids, and other health-promoting phytochemicals [1,2]. However, global citrus production faces significant challenges from diseases like Huanglongbing (HLB), caused by *Candidatus Liberibacter asiaticus* (CLas), and citrus canker, caused by *Xanthomonas citri* subsp. citri. These diseases greatly reduce tree productivity, fruit quality, and orchard longevity worldwide [3,4,5]. These challenges demand the development of new citrus cultivars with improved disease resistance and key agronomic traits. Consequently, breeders have long worked to develop improved scion and rootstock cultivars through traditional breeding methods. However, these methods are time-consuming and labor-intensive and can take around 20 years from a cross to cultivar release [6]. Additionally, the complex genome structure and high heterozygosity of cultivated citrus [7], together with nucellar polyembryony, self-incompatibility, partial sterility, and a 5–8-year juvenile phase [8], make citrus breeding especially challenging. Therefore, there is a need to integrate modern precision breeding tools with conventional approaches to increase genetic gain per breeding cycle.

However, the successful application of these technologies depends on efficient, reproducible plant regeneration and transformation. Among gene delivery methods, *Agrobacterium*-mediated transformation is the most common in citrus because of its simplicity, broad applicability, and lack of requirement for protoplast culture or embryogenic cell suspensions [9, 10]. Despite continuous improvements, transformation efficiency remains highly genotype-dependent and is often limited by prolonged *in vitro* culture, low recovery of transgenic events, and the occurrence of escapes and chimeric regenerants under antibiotic selection [11,12,13]. Most protocols have been optimized predominantly using juvenile seedling explants, especially epicotyl, cotyledon, and hypocotyl segments via direct organogenesis, whereas indirect organogenesis has been reported for fewer genotypes [11,14]. Indirect organogenesis involves a prolonged callus proliferation phase prior to shoot regeneration, providing an extended window during which transformed cells can proliferate and be selected, potentially improving the recovery of stable transgenic regenerants [15,16]. However, efficient and reproducible transformation systems based on indirect organogenesis remain limited in citrus.

Establishing an efficient regeneration and transformation system alone is insufficient for successful citrus transformation; reliable identification of transformed tissues throughout the prolonged regeneration process is equally important. Although selectable marker genes favorably select transformed cells, antibiotic selection alone does not always eliminate non-transformed tissues, leading to escapes and chimeric regenerants during prolonged culture [10,17]. Therefore, reporter genes are routinely incorporated into transformation vectors to facilitate identification and recovery of transgenic events. The most widely used reporters are protein-based, including β-glucuronidase (*GUS*), luciferase (*LUC*), and green fluorescent protein (*GFP*), each with inherent limitations for long-term regeneration studies. *GUS* requires destructive histochemical staining, precluding repeated assessment of the same explant [18]. *LUC* depends on exogenous substrate application and specialized imaging. *GFP* enables non-destructive, real-time visualization and has been widely used in citrus transformation [19], but its detection requires fluorescence imaging equipment and may be complicated by chlorophyll autofluorescence in regenerating green tissues [20,21,22]. These limitations have spurred the development of visible pigment reporters, which produce colored metabolites that can be detected directly under ambient light without exogenous substrates or specialized instrumentation [20,23].

Visible pigment reporters can be broadly divided into anthocyanin- and betalain-based systems, which differ fundamentally in their mechanisms of pigment production. *ROSEA1* is an R2R3-MYB transcription factor that activates the endogenous anthocyanin biosynthetic pathway through host regulatory networks [24], whereas the *RUBY* reporter encodes the three enzymes required for betalain biosynthesis as a single polycistronic cassette, enabling pigment production largely independent of endogenous anthocyanin regulatory pathways [20]. Both systems have been successfully applied for visual selection of transformed tissues in diverse plant species, including woody horticultural crops such as Mexican lime, bilberry, and apple [25,26,27]. However, these reporters have primarily been evaluated independently, and their performance during the successive stages of regeneration has not been systematically investigated in citrus, particularly in transformation systems based on indirect organogenesis.

In this study, an optimized *Agrobacterium*-mediated transformation system based on indirect organogenesis was established for Carrizo citrange. Using a common *GFP–NPTII* backbone, the anthocyanin reporter *ROSEA1 (ROS1)* and the betalain reporter *RUBY* were evaluated as visible markers throughout the transformation process, from callus induction to transgenic plant recovery. The primary objective was to establish and validate an indirect organogenesis-based transformation workflow for Carrizo citrange and to determine whether visible pigmentation could provide continuous, nondestructive tracking of transformed tissues throughout regeneration, using *GFP* as a common reference.

## 2. Results

### 2.1 Establishment of an indirect organogenesis-based transformation system for Carrizo citrange using visible reporter markers

An *Agrobacterium*-mediated indirect organogenesis system was developed for Carrizo citrange to evaluate anthocyanin- and betalain-producing genes as visible reporters of transformation (Fig. 1). Three constructs were assembled: the unmodified *pOX135* vector as a control (CK), and *pOX135*-*ROSEA1* and *pOX135-RUBY*, in which anthocyanin and betalain biosynthesis, respectively, are driven by a 2×CaMV35S promoter (Fig. 1c). All three contain the fused *eGFP– NPTII* cassette, so that each visible marker can be evaluated against *GFP* within a common vector background. Correct assembly of all constructs was confirmed by PCR prior to transformation (Fig. 1d).

**Figure 1.**
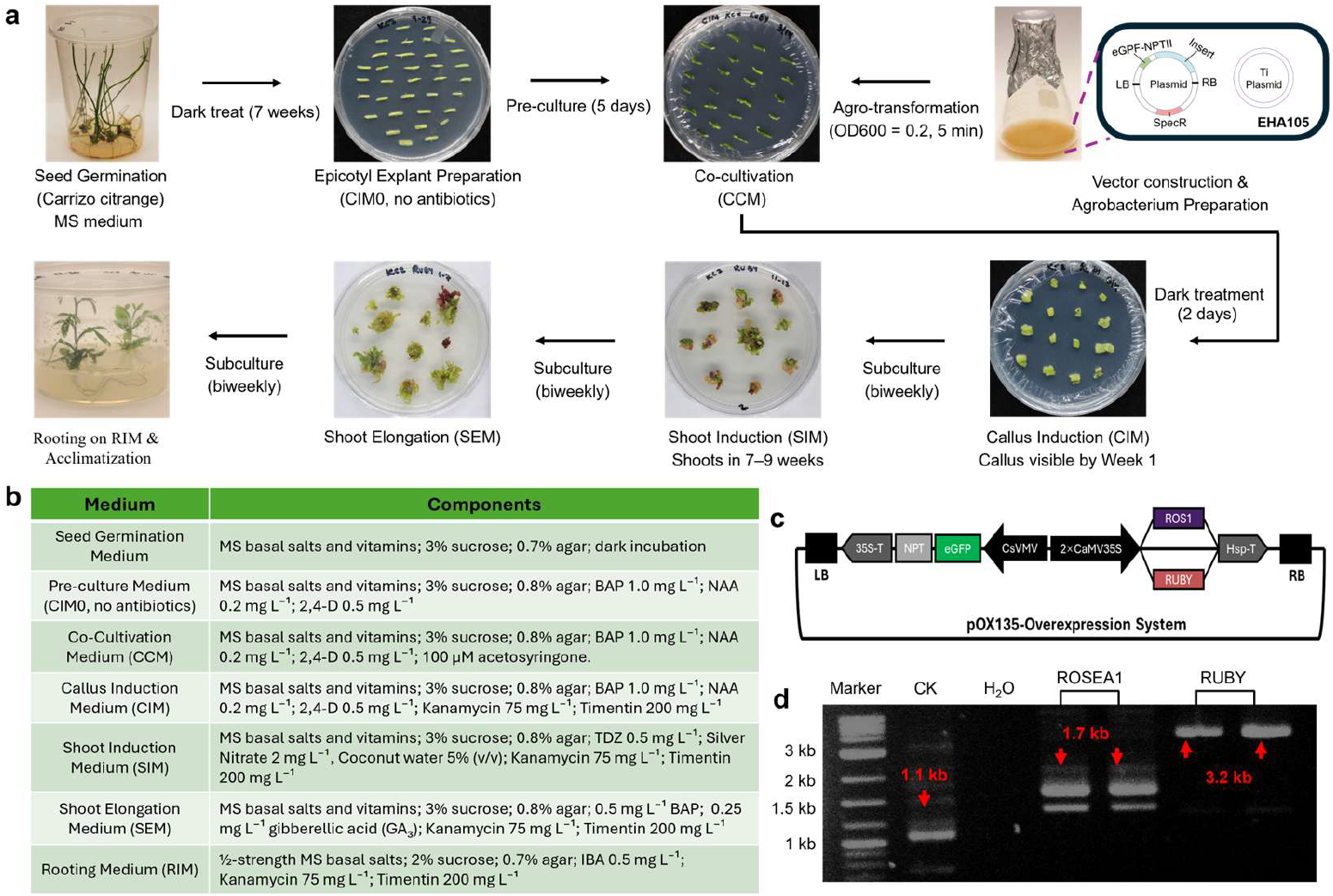
Establishment of an indirect organogenesis-based citrus transformation system using visible reporter markers. **(a)** Overview of the transformation and regeneration workflow, including seed germination, epicotyl preparation, pre-culture, *Agrobacterium* infection and co-cultivation, callus induction and selection, shoot regeneration and elongation, and rooting. **(b)** Composition of the culture media used at each stage of transformation and regeneration. CIM, callus induction medium; CCM, co-cultivation medium; SIM, shoot induction medium; SEM, shoot elongation medium; RIM, rooting medium. **(c)** Simplified T-DNA structures of the pOX135-based constructs. The control construct (CK) contains the CsVMV-driven *eGFP*-NPT*II* reporter-selection fusion cassette, whereas pOX135-*ROSEA1* (*ROS1*) and pOX135-*RUBY* additionally contain the visible reporter *ROSEA1* or *RUBY*, respectively, driven by the 2×CaMV 35S promoter and terminated by Hsp-T. **(d)** Diagnostic PCR of the pOX135-derived plasmid constructs. Expected amplicons were 1.1 kb for CK, 1.7 kb for pOX135-*ROSEA1* (*ROS1*), and 3.2 kb for pOX135-*RUBY*. M, DNA marker; H_2_O, no-template control.

Etiolated epicotyl segments excised from dark-grown seedlings were used as explants and were carried through preculture, *Agrobacterium* inoculation, and co-cultivation, followed by callus induction, shoot induction, shoot elongation, and rooting under continued kanamycin selection (Fig. 1a). Regenerated shoots were then transferred to a rooting medium containing kanamycin 75 mg L^-1^ and Timentin 200 mg L^-1^. Media compositions for each stage are given in Fig. 1b. Callus became visible within one week of transfer to selection medium, early shoot regeneration was detectable by week 5, with clearly developed adventitious shoots becoming prominent during weeks 7–9, and rooted plantlets were obtained by week 20.

### 2.2. Visible reporter expression enables non-destructive identification of transformed callus during induction and proliferation

To determine whether constitutive visible reporter expression affected dedifferentiation, callus formation was compared among CK, *ROSEA1*, and *RUBY* explants over seven weeks on selective medium. Visible callus initiated at the cut surfaces of epicotyl explants within 7 days in all three treatments (Fig. 2a). Control explants formed pale-yellow callus, whereas *ROSEA1* and *RUBY* tissues developed distinct pink-purple and carmine-red pigmentation, respectively, from week 3; pigmentation intensified by week 7, allowing reporter-expressing callus to be distinguished under ambient light (Fig. 2a). Equivalent phenotypes were observed across independent biological replicates (Fig. S1–S3).

**Figure 2.**
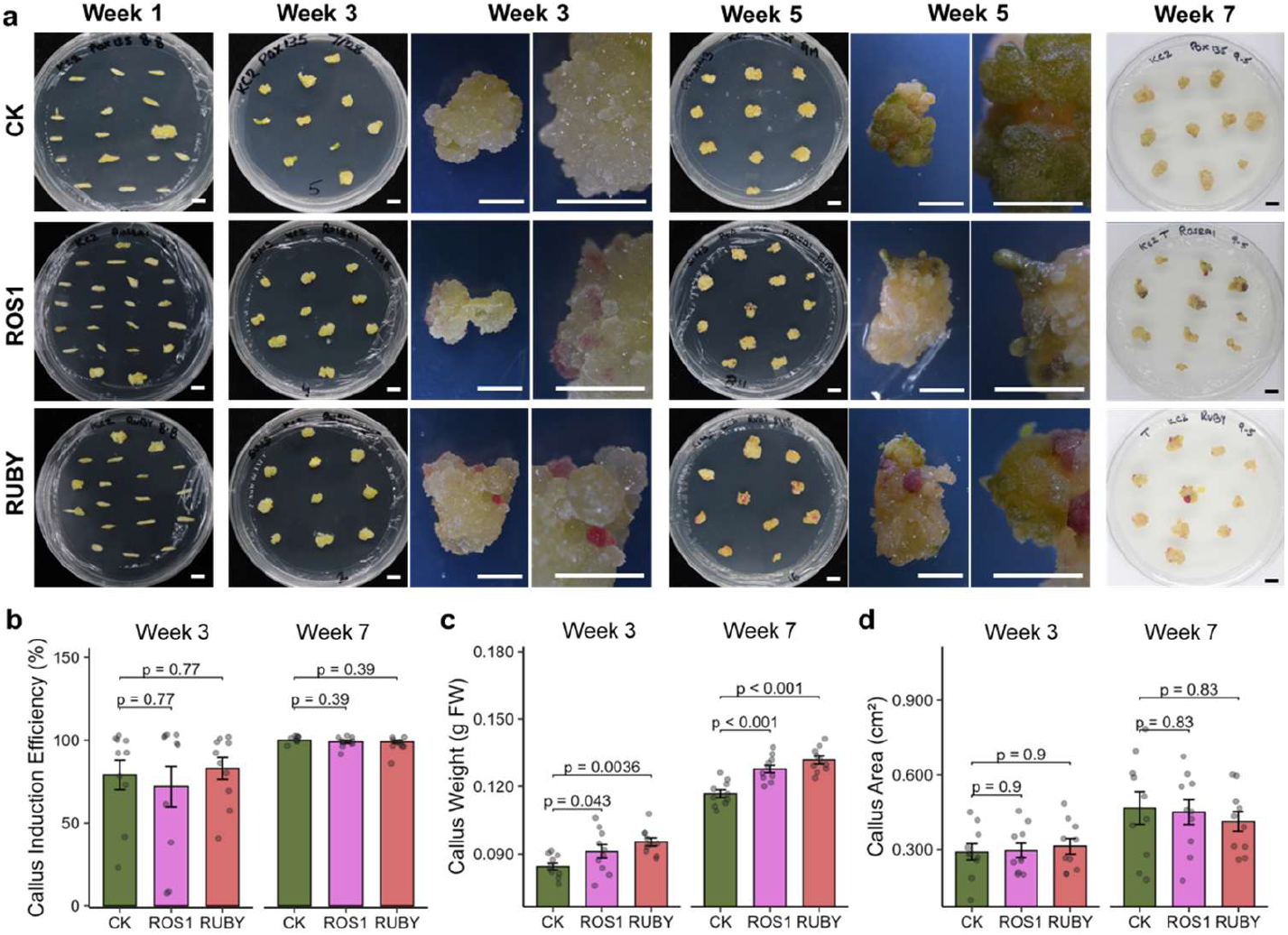
Visible markers *ROSEA1* and *RUBY* indicate callus growth status during transformation in citrus. **(a)** Representative phenotypes of citrus explants during *Agrobacterium*-mediated transformation under control (CK), *ROSEA1* (*ROS1*), and *RUBY* treatments at weeks 1, 3, 5, and 7. CK explants developed original calli without visible pigment accumulation, whereas *ROSEA1* and *RUBY* explants displayed pink-purple anthocyanin and betalain pigmentation, respectively. Representative Petri-dish overviews and close-up views of individual calli are shown. **(b)** Callus induction efficiency (%) **(c)** Callus weight (g fresh weight) **(d)** Callus area (cm^2^). CK (green), *ROSEA1* (purple), and *RUBY* (carmine) treatments at weeks 3 and 7. Data are presented as mean ± s.d. (n = 8–10 biological replicates). Statistical significance was assessed using two-tailed Student’s t-tests (CK versus *ROSEA1* ; CK versus *RUBY*). Scale bars, 1 cm (Petri dishes in a) and 5 mm (close-ups in a).

Callus induction efficiency was unaffected by either reporter. Approximately 70–80% of explants formed callus by week 3 and close to 100% by week 7, with no significant difference between either marker treatment and CK at either timepoint (Fig. 2b; week 3, p = 0.77; week 7, p = 0.39). In contrast, callus fresh weight was significantly higher in both reporter treatments than in CK. At week 3, *ROSEA1* and *RUBY* calli were heavier than CK (p = 0.043 and p = 0.0036, respectively), and this difference became more pronounced by week 7, when both markers exceeded CK at p < 0.001 (Fig. 2c). Callus area, however, did not differ significantly among treatments at either timepoint (Fig. 2d; week 3, p = 0.9; week 7, p = 0.83), indicating that the increase in biomass was not accompanied by greater lateral expansion of the callus. Overall, callus induction remained robust across all three constructs, while reporter-specific pigmentation enabled transformed callus to be readily distinguished during proliferation.

### 2.3. Visible pigmentation enables continuous visual tracking of transformed tissues during shoot regeneration

Following callus induction, shoot regeneration was compared among CK, *ROSEA1*, and *RUBY* transformed explants on shoot induction medium, and regeneration efficiency and shoot number were scored at weeks 5, 7, and 9. Adventitious shoots developed from callus in all three treatments, exhibiting pink-purple pigmentation in *ROSEA1* tissues and carmine-red in *RUBY* tissues, which also appeared in the developing shoots (Fig. 3a). However, CK shoots showed no pigmentation (Fig. 3a). GFP fluorescence verified transgene expression in regenerating tissues across all three treatments.

**Figure 3.**
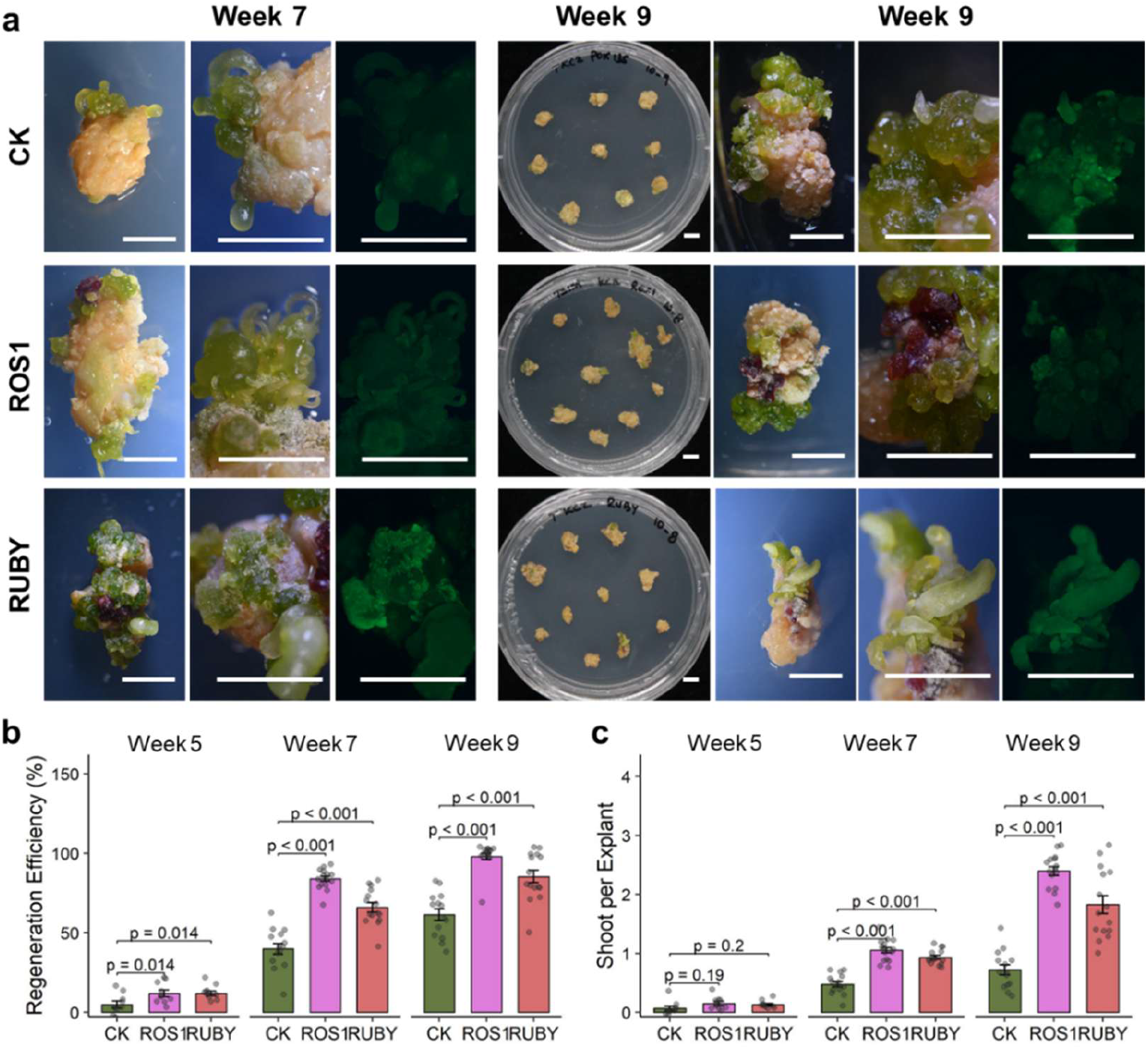
Visible markers *ROSEA1* and *RUBY* indicate plant regeneration status during transformation in citrus. **(a)** Representative phenotypes of citrus explants during *Agrobacterium*-mediated transformation under control (CK), *ROSEA1* (*ROS1*), and *RUBY* treatments at weeks 7 and 9. CK explants showed original primordia without visible pigment accumulation, whereas *ROSEA1* and *RUBY* explants displayed pink-purple anthocyanin and betalain pigmentation, respectively, with colored shoot formation. Representative Petri-dish overviews, close-up views of individual explants, and GFP fluorescence images are shown. **(b)** Regeneration efficiency (%) **(c)** Shoots per explant in CK (green), *ROSEA1* (purple), and *RUBY* (carmine) treatments at weeks 5, 7, and 9. Data are presented as mean ± s.d. (n = 8–10 biological replicates). Statistical significance was assessed using two-tailed Student’s t-tests (CK versus *ROSEA1*; CK versus *RUBY*). Scale bars, 1 cm (Petri dishes in a) and 5 mm (close-ups in a).

Regeneration efficiency remained relatively low at week 5 but was higher in both reporter treatments than in CK (Fig. 3b; Fig. S2). These differences became substantially more pronounced at weeks 7 and 9, when both *ROSEA1* and *RUBY* showed markedly higher regeneration efficiencies than CK. Regeneration efficiency in *ROSEA1* and *RUBY* reached approximately 82% and 65% at week 7 and 90% and 82% at week 9, compared with 40% and 62% in CK (p < 0.001 for both markers at both timepoints; Fig. 3b). Shoot number followed the same pattern. Shoots per explant did not differ significantly among treatments at week 5 (p = 0.19–0.2), but by week 7 both markers exceeded CK (p < 0.001), and by week 9 *ROSEA1* and *RUBY* produced approximately 2.4 and 1.8 shoots per explant, respectively, compared with 0.75 in CK (p < 0.001; Fig. 3c; Fig. S4). Overall, both reporter constructs remained compatible with efficient shoot regeneration, while visible pigmentation allowed transformed tissues to be continuously followed during the transition from callus to shoots.

### 2.4. Visible pigmentation provides an alternative to GFP fluorescence for identifying transformed callus

After characterizing reporter visibility and regeneration phenotypes at the callus and shoot stages, we next quantitatively compared visible pigmentation with GFP-based detection, beginning with the callus stage. Since all three constructs carry the same *eGFP*–*NPTII* cassette, GFP fluorescence served as a common benchmark for evaluating reporter performance. Transformation efficiency was assessed by GFP fluorescence and, for the visible reporter lines, by pigmentation at weeks 3 and 7. Transformed calli showed GFP fluorescence across all treatments, while *ROSEA1* and *RUBY* calli additionally developed pink-purple and carmine-red pigmentation, respectively. In contrast, non-transformed calli showed neither GFP fluorescence nor visible pigmentation (Fig. 4a).

**Figure 4.**
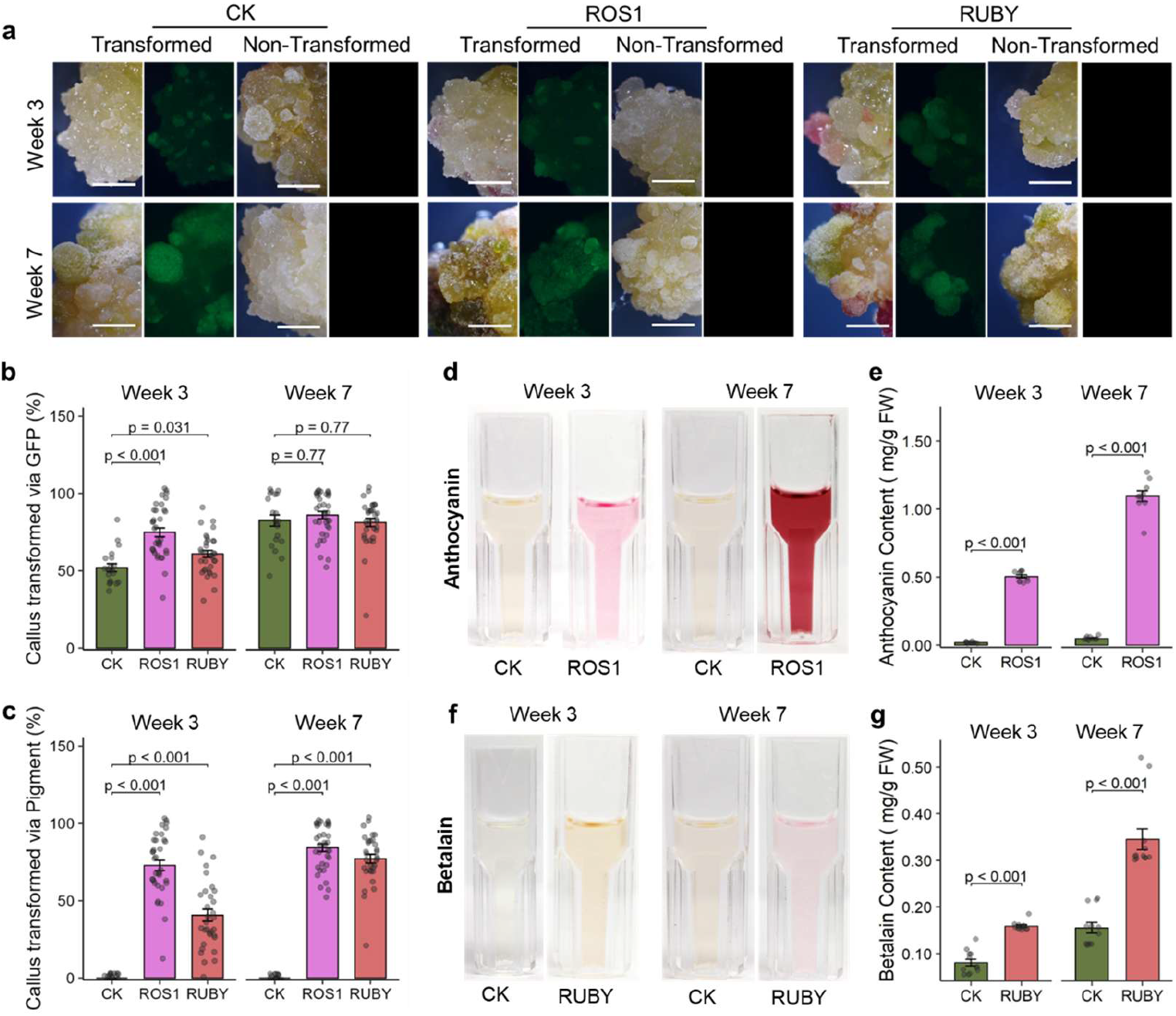
Visible markers *ROSEA1* and *RUBY* indicate callus transformation status in citrus. **(a)** Representative transformed and non-transformed citrus calli during *Agrobacterium*-mediated transformation under control (CK), *ROSEA1* (*ROS1*), and *RUBY* treatments at weeks 3 and 7. Reporter-positive calli showed GFP fluorescence across all three treatments, whereas *ROSEA1*- and *RUBY*- containing calli additionally displayed visible pigmentation. Non-transformed calli lacked GFP fluorescence and visible reporter pigmentation. Scale bar, 5 mm. **(b, c)** Callus transformation efficiency indicated via GFP fluorescence (%) (b) and visible pigmentation (%) (c) in CK (green), *ROSEA1* (purple), and *RUBY* (carmine) treatments at weeks 3 and 7. **(d, e)**Representative anthocyanin extraction (d) and anthocyanin content (mg/g FW) (e) in CK and *ROSEA1* treatments at weeks 3 and 7. **(f, g)**Representative betalain extraction (f) and betalain content (mg/g FW) (g) in CK and *RUBY* treatments at weeks 3 and 7. Data are presented as mean ± s.d. (n = 8–10 biological replicates). Statistical significance was assessed using two-tailed Student’s t-tests (CK versus *ROSEA1*; CK versus *RUBY*).

GFP-based transformation efficiency varied among treatments at week 3 but increased across all three constructs during continued culture (Fig. 4b). By week 7, GFP-based efficiencies had converged to approximately 82–85%, with no significant differences among treatments (p = 0.77). Thus, despite differences observed earlier in culture, all three constructs reached comparable GFP-based transformation frequencies by week 7.

Pigment-based scoring detected no positive calli in CK at either timepoint (Fig. 4c). In *ROSEA1*, pigment-based detection was already high at week 3 (approximately 74%) and increased to approximately 85% by week 7. *RUBY* pigmentation was detected in a smaller proportion of calli at week 3 (approximately 40%) but increased to approximately 78% by week 7 (p < 0.001 versus CK at both timepoints). By week 7, pigment-based and GFP-based detection had converged to comparable levels within each reporter line. In contrast, at week 3, *RUBY* pigmentation lagged behind both its corresponding *GFP* signal and *ROSEA1* pigmentation, indicating a slower onset of visually detectable betalain accumulation.

To confirm that the visible signals indicated true pigment accumulation, pigments were extracted from callus tissue and quantified. Anthocyanin content in *ROSEA1* was notably higher than in CK at both timepoints and increased markedly between weeks 3 and 7 (Fig. 4d,e; p < 0.001). Similarly, betalain levels in *RUBY* exceeded those in CK and increased over the same period (Fig. 4f,g; p < 0.001). This gradual accumulation correlated with the increased detection of pigments from week 3 to 7, establishing a direct link between the visible signals and actual pigment content.

### 2.5. ROSEA1 and RUBY show distinct visual detectability during shoot regeneration

To determine whether visible pigmentation indicates transformation status in regenerating shoots, where chlorophyll accumulation and tissue differentiation might affect marker visibility compared to callus, transformation was evaluated by GFP fluorescence and visible pigmentation at weeks 13 and 17 (Fig. 5a), with week 15 intermediate shoot-stage (Fig. S5). *ROSEA1* and *RUBY* shoots showed pink-purple and carmine-red pigmentation, respectively, while CK shoots remained unpigmented. In all three treatments, GFP fluorescence confirmed transformation.

**Figure 5.**
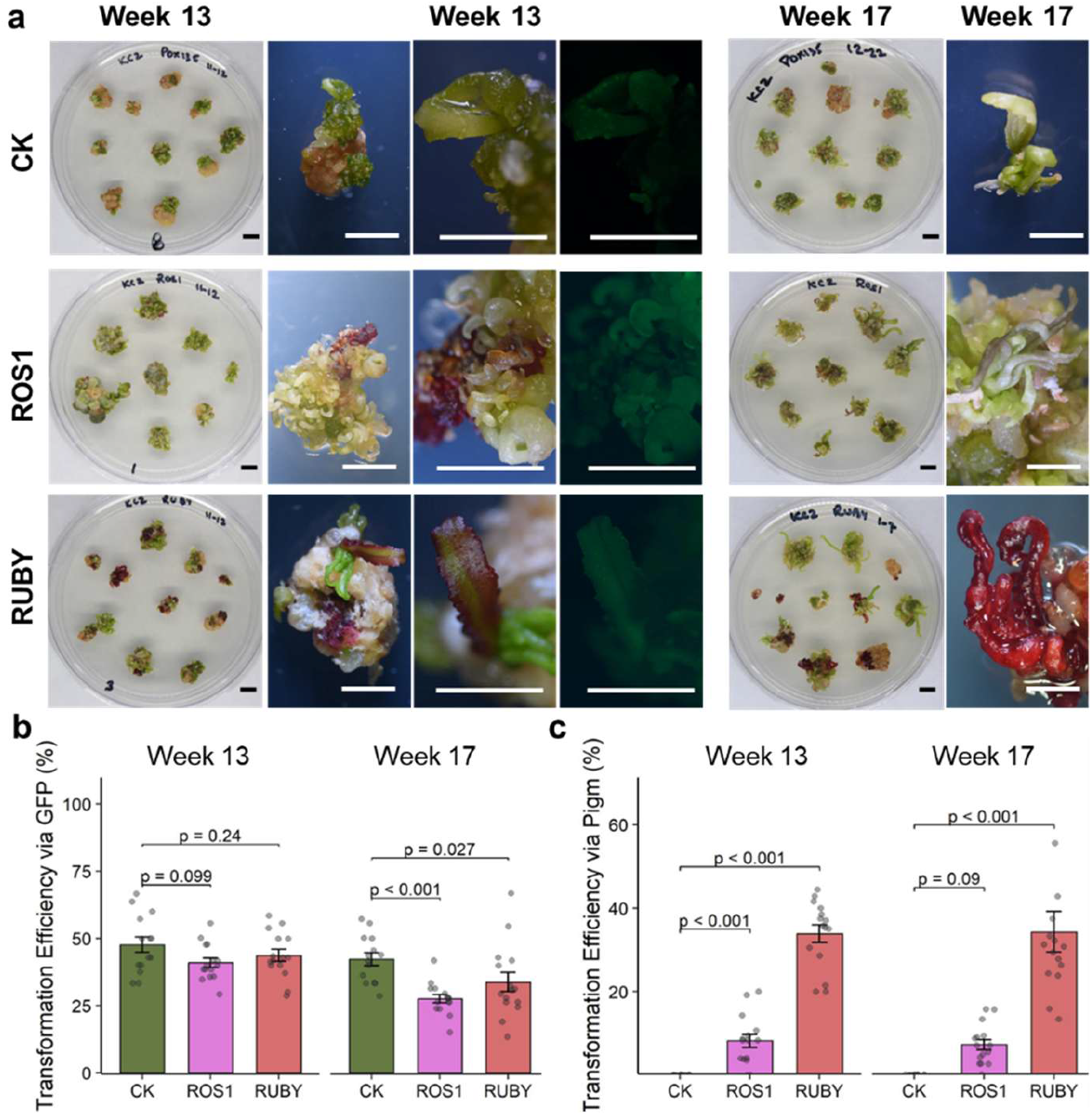
Visible markers *ROSEA1* and *RUBY* indicate shoot transformation status in citrus. **(a)** Representative phenotypes of citrus shoot development during *Agrobacterium*-mediated transformation under control (CK), *ROSEA1* (*ROS1*), and *RUBY* treatments at weeks 13 and 17. CK shoots developed without visible pigment accumulation, whereas *ROSEA1* and RUBY shoots displayed pinkpurple anthocyanin and betalain pigmentation, respectively. Representative Petri-dish overviews, close-up views of individual shoots, and GFP fluorescence images are shown. **(b, c)** Shoot transformation efficiency indicated via GFP fluorescence (%) (b) and visible pigmentation (%) (c) in CK (green), *ROSEA1* (purple), and *RUBY* (carmine) treatments at weeks 13 and 17. Data are presented as mean ± s.d. (n = 8–10 biological replicates). Statistical significance was assessed using two-tailed Student’s t-tests (CK versus *ROSEA1*; CK versus *RUBY*). Scale bars, 1 cm (Petri dishes in a) and 5 mm (close-ups in a).

GFP-based transformation efficiency was comparable among the three treatments at week 13 (approximately 42–48%; Fig. 5b; p = 0.099 and p = 0.24 for *ROSEA1* and *RUBY* versus CK) but declined by week 17, when both marker lines were significantly lower than CK (*ROSEA1*, p < 0.001; *RUBY*, p = 0.027). This decline in GFP-based efficiency occurred despite all constructs carrying the same *eGFP–NPTII* cassette.

Pigment-based scoring produced no positive shoots in CK at either timepoint, confirming marker specificity, but the two pigment markers diverged markedly (Fig. 5c). *RUBY*-based detection reached approximately 34% at both timepoints (p < 0.001 versus CK), comparable to the GFP-based efficiency in the same line by week 17. *ROSEA1*-based detection, by contrast, remained low throughout, at approximately 8% at week 13 and 7% at week 17, and was significantly above CK only at week 13 (p < 0.001), with the difference no longer significant by week 17 (p = 0.09). Anthocyanin pigmentation therefore identified only a small proportion of transformed shoots at this stage, well below the corresponding GFP-based efficiency. These results indicate that, in contrast to the callus stage, at which both markers reported transformation efficiently, only *RUBY* provided visual detection of transformed shoots comparable to *GFP*, whereas *ROSEA1*-based detection was substantially reduced during shoot regeneration.

### 2.6. Recovery and molecular confirmation of stable transgenic plantlets

Rooted plantlets were recovered from all three treatments and developed into fully formed plantlets by week 20 (Fig. 6a, Fig. S6). GFP fluorescence was detected in the leaves and root tips of plantlets from all three treatments, consistent with stable expression of the shared *eGFP–NPTII* cassette in differentiated tissues (Fig. 6a). Visible pigmentation was observed in the regenerated *ROSEA1* and *RUBY* tissues but not in CK (Fig. 6a).

**Figure 6.**
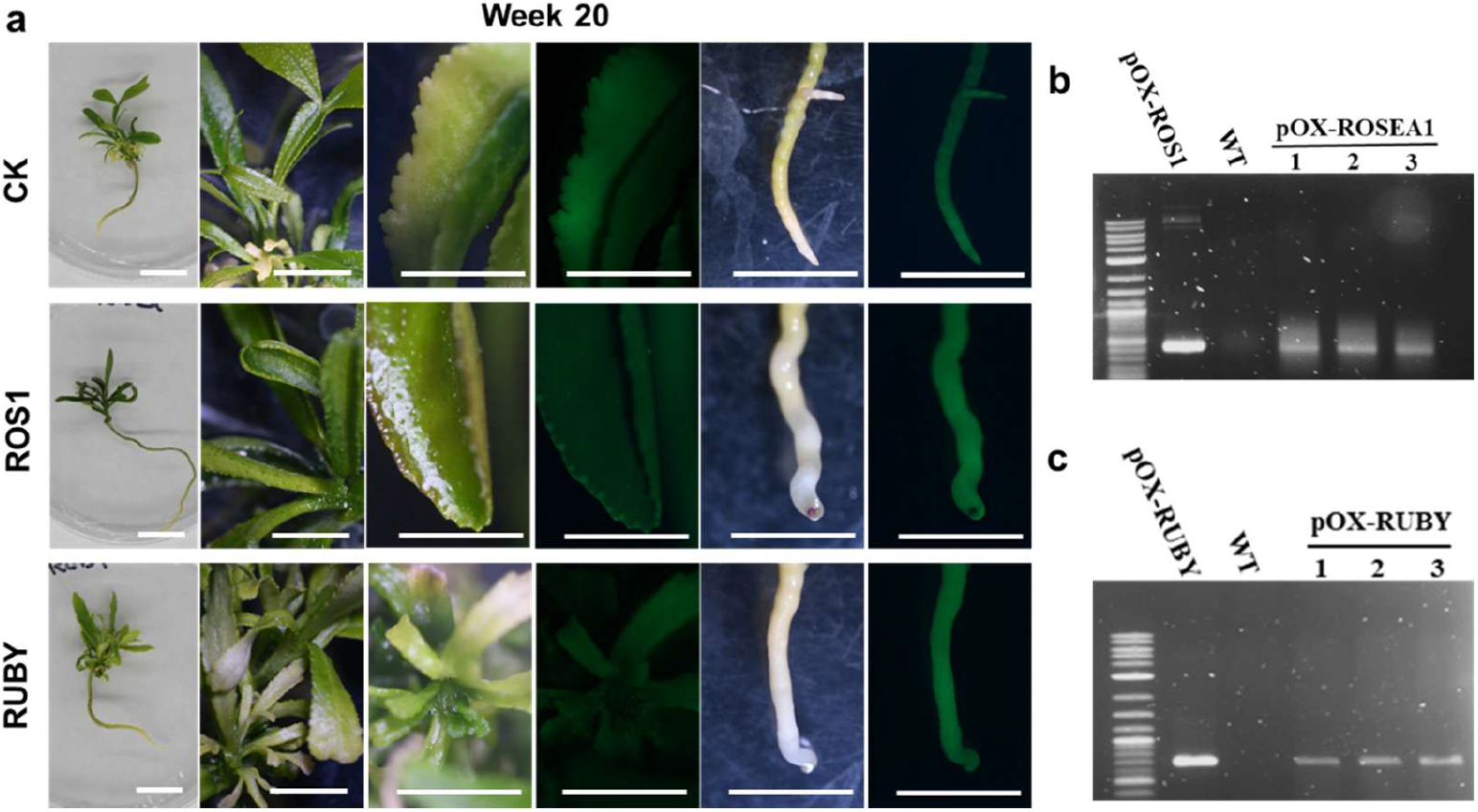
Transgenic citrus plantlets stably expressing *ROSEA1* and *RUBY* visible markers at week 20. **(a)** Representative fully developed transgenic citrus plantlets under control (CK), *ROSEA1* (*ROS1*), and *RUBY* treatments at week 20. Overview images, close-up views, and GFP fluorescence detection in leaves and root tips are shown. PCR genotyping confirmation of *ROSEA1*- overexpressing **(b)** and *RUBY*-overexpressing **(c)** transgenic lines. pOX-*ROSEA1* and pOX-*RUBY* vectors served as positive controls, wild-type (WT) as negative control, and three independent transgenic lines (1–3) are shown. Scale bars, 1 cm (whole plantlets in a) and 5 mm (close-ups in a).

The presence of the transgenes in regenerated plantlets was confirmed by PCR genotyping of genomic DNA *from* independent regenerated lines (Fig. 6b,c). Amplification with *ROSEA1*- and *RUBY*-specific primers yielded products of the expected size in three independent pOX-*ROSEA1* lines and three independent pOX-*RUBY* lines, respectively, corresponding to the amplicons obtained from the positive plasmid controls, whereas no product was amplified from wildtype genomic DNA in either assay (Fig. 6b,c). Taken together with the GFP fluorescence detected in leaves and root tips, these results confirm the recovery of stably transformed citrus plantlets carrying the *ROSEA1* or *RUBY* visible marker.

## 3. Discussion

Citrus is widely regarded as recalcitrant to *Agrobacterium*-mediated transformation, with success rates ranging from 0 to about 45% and strongly dependent on factors such as species, genotype, explant type, and selection strategy [10,12,19,28]. Moreover, in citrus, a major constraint is not the transformation process itself but the reliable identification of transformed tissue, because during the prolonged regeneration, antibiotic selection may permit non-transformed escapes and chimeric shoots to persist [9,17]. Scorable reporter genes that permit visual identification of transformed tissue offer a direct means of addressing this bottleneck [20,29]. Here, we established an indirect organogenesis-based transformation workflow for Carrizo citrange and evaluated *ROSEA1* and *RUBY* as visible reporters for tracking transformed tissues throughout the regeneration process using a shared *GFP*–*NPTII* marker as a reference. Thus, the major practical contributions of this study are the development of an indirect regeneration route for citrus transformation and the integration of non-destructive visual monitoring throughout this prolonged workflow.

### 3.1. An efficient indirect organogenesis-based transformation system

Efficient recovery of transgenic citrus plants remains challenging and requires effective gene delivery and robust regeneration systems. Although advances have been made in *Agrobacterium*-mediated transformation through optimization of bacterial strains, infection, and co-cultivation [13,30], most protocols produce shoots directly from juvenile epicotyl explants. Few studies have developed transformation systems based on indirect organogenesis [31]. This study presents an efficient indirect organogenesis-based citrus transformation system using juvenile tissue from Carrizo citrange that supported rapid callus establishment, efficient shoot regeneration, and direct recovery of rooted transgenic plantlets. Callus developed within one week after transfer to selection medium, and regenerated shoots were obtained within nine weeks, allowing complete recovery of plantlets in approximately 20 weeks.

A key distinction between the two regeneration pathways is the cellular origin of regenerated shoots. In direct organogenesis, adventitious shoots develop directly from wounded explant tissues, often involving multiple neighboring competent cells, which increases the likelihood of recovering chimeric regenerants after *Agrobacterium*-mediated transformation [10,17,32,33]. In contrast, indirect organogenesis includes a proliferative callus phase before shoot induction, allowing transformed cells to expand before organogenic commitment and thereby providing a biological basis for recovering more uniformly transformed regenerants [16,34]. Although chimerism was not directly assessed in the present study, the generally uniform reporter expression observed throughout regenerated calli and shoots is consistent with regeneration from predominantly transformed cell populations. Future histological or single-cell approaches will be required to determine whether indirect organogenesis consistently reduces chimerism compared to direct regeneration in citrus.

A further practical advantage of the protocol presented in this study was the recovery of rooted shoots without shoot-tip micrografting. Because regenerated citrus shoots often exhibit poor adventitious rooting, many transformation protocols depend on micrografting onto seedling rootstocks to establish transgenic plants [11,21,35]. In contrast, in the present study, regenerated shoots rooted readily on IBA-containing medium without micrografting, thereby simplifying the transformation workflow and reducing both labor and recovery time. Combined with visible reporter-based monitoring, the indirect organogenesis system described here expands the regeneration strategies available for citrus transformation, including applications in genome editing and functional genomics that require sustained tissue proliferation and reliable identification of transformed tissues over extended culture periods.

### 3.2. Visible reporters are compatible with callus proliferation

Callus induction was unaffected by either reporter: callus induction efficiency and callus area were equivalent across treatments, and callus formed within roughly one week of transfer to selection medium. Callus fresh weight, however, was significantly greater in both reporter lines than in the control at both timepoints, while callus area was unchanged, indicating greater biomass per unit area in the pigmented lines. Importantly for the intended application of the reporters, neither *ROSEA1* nor *RUBY* compromised callus establishment or proliferation, while their pigmentation allowed transformed callus to be readily distinguished during this stage of the workflow.

The similar biomass response observed in *ROSEA1*- and *RUBY*-expressing calli may reflect physiological differences associated with pigment accumulation, although this was not a primary objective of the present study. Anthocyanins and betalains are functionally convergent metabolites that both scavenge reactive oxygen species (ROS) and act as osmotic regulators, thereby protecting cells from oxidative and osmotic stress [36,37,38]. During transformation, explants are exposed to multiple sources of stress, including wounding, *Agrobacterium* infection, and prolonged *in vitro* culture, all of which are associated with enhanced ROS production and oxidative stress that can limit cell proliferation and regeneration [39,40,41]. Although these properties provide a possible context for the greater fresh weight observed in the reporter treatments, the present data do not establish a mechanistic relationship between pigment accumulation and callus biomass.

Importantly, the enhanced callus biomass observed in the reporter lines did not correspond with any detectable reduction in callus proliferation. This finding is noteworthy because constitutive anthocyanin accumulation has previously been associated with growth inhibition in citrus. Mexican lime plants overexpressing the anthocyanin regulator *RUBY* exhibited dwarfism and leaf curling, indicating that excessive pigment accumulation can impose developmental penalties [25]. Similar growth trade-offs have also been reported in other species with elevated anthocyanin accumulation, where strong activation of the flavonoid pathway has been associated with growth and developmental defects in heterologous hosts [42]. In contrast, neither *ROSEA1* nor *RUBY* expression adversely affected callus growth in the present study, suggesting that pigment accumulation remained within a physiologically tolerated range during the callus stage.

To the best of our knowledge, previous studies using *ROSEA1*- or *RUBY*-based visible reporters have focused primarily on reporter performance during transformation and regeneration for tracking of transformation and have not evaluated their effects on callus biomass [23,43,44]. The biomass difference observed here should therefore be considered a secondary observation rather than evidence that either reporter promotes callus growth. Overall, these findings support the use of *ROSEA1* and *RUBY* as visual markers for citrus transformation, with no apparent negative effect on callus proliferation.

### 3.3. Reporter expression is compatible with shoot regeneration

Both reporter constructs remained compatible with efficient shoot regeneration throughout the Carrizo indirect organogenesis workflow. Although regeneration efficiency and shoot number were higher in the reporter treatments than in the control from week 7 onward, these differences were not a primary endpoint of the study and should not be interpreted as evidence that either reporter promotes regeneration. This distinction is relevant because constitutive pigment-marker expression has been reported to hinder regeneration in both anthocyanin- and betalain-based systems in some species. Constitutive *ROSEA1* expression reduced regeneration in several eudicot species [45], whereas *RUBY*-expressing bilberry calli showed reduced recovery of pigmented shoots during callus-mediated regeneration [26]. The absence of an obvious regeneration penalty in Carrizo therefore indicates that both visible reporters can be incorporated into this indirect organogenesis system without preventing efficient shoot recovery.

Several features of the experimental system may contribute to the difference from previous reports, including regeneration route, hormonal environment, species, and selection regime. The *ROSEA1*-associated regeneration penalty reported previously was most pronounced during direct organogenesis and stable overexpression, whereas altered cytokinin availability influenced this response [45]. In the present study, Carrizo regenerated through a proliferative callus phase under cytokinin-rich conditions before shoot induction. However, these factors were not independently tested here, and the present study was not designed to identify the biological basis of the differences in regeneration among treatments.

A second consideration is that the higher regeneration observed in the reporter treatments may partly reflect preferential recovery of transformed, regeneration-competent tissues during continuous selection rather than a direct developmental effect of reporter expression. In citrus, non-transformed cells may persist within or adjacent to transformed tissues under kanamycin selection, contributing to escapes and chimeric regeneration [17,33]. Visible pigmentation may therefore facilitate identification of genuinely transformed sectors during repeated subculture. Accordingly, the relevant conclusion for the present methodology is that *ROSEA1* and *RUBY* were compatible with efficient shoot formation and provided visible tracking of transformed tissues during the callus-to-shoot transition, rather than that either reporter directly enhanced regeneration.

### 3.4. Anthocyanin and betalain reporters exhibit distinct performance across developmental stages

This study integrated visible reporters into an indirect organogenesis-based citrus transformation system, enabling continuous visual tracking of transformed tissues during regeneration. At the callus stage, pigmentation closely corresponded with GFP fluorescence, while control tissues showed no reporter-specific pigmentation, supporting the specificity of both visible reporter systems during early regeneration. At the plantlet stage, continued GFP fluorescence together with diagnostic PCR confirmation of independent regenerated lines supported that visible pigmentation in recovered plantlets reflected stable transformation rather than solely transient expression. These findings demonstrate that visible pigmentation can complement fluorescence-based screening for monitoring transformed tissues throughout the citrus regeneration process. This stage-by-stage validation is particularly relevant for citrus, where prolonged regeneration and repeated subculture make continuous identification of transformed tissues important for reliable plant recovery.

Although both reporters showed similar performance during callus proliferation, they diverged after shoot induction. *RUBY* maintained robust, consistent pigmentation during shoot regeneration, while *ROSEA1*-associated pigmentation declined despite continued GFP fluorescence and PCR confirmation. This pattern is consistent with developmental regulation of pigment accumulation rather than being readily explained by transgene loss. Thus, reporter performance should be evaluated across the complete regeneration process rather than inferred from a single developmental stage.

The contrasting behavior of the two reporters is consistent with their underlying biology. *ROSEA1* functions by activating the endogenous anthocyanin biosynthetic pathway through the MYB–bHLH–WD40 regulatory complex, whose activity depends on tissue identity, developmental stage, light, and the availability of endogenous regulatory partners [24,46]. Consequently, anthocyanin accumulation frequently varies during development and among tissues. By contrast, *RUBY* encodes the complete betalain biosynthetic pathway as a single polycistronic cassette and synthesizes pigment independently of endogenous transcriptional regulators [20]. This difference may contribute to the more persistent *RUBY* pigmentation observed during shoot regeneration. *RUBY* has been successfully applied as a visible reporter in diverse plant systems [20,29]. Similarly, studies of anthocyanin-engineered citrus have reported stronger pigmentation in juvenile tissues than in mature tissues, supporting the broader observation that anthocyanin-based visual signals can vary substantially with developmental stage [47–49].

The observed decline in GFP-based detection during the shoot stage indicates that the proportion of regenerated shoots with detectable *GFP* decreased during prolonged culture. This pattern may reflect factors such as transgene silencing, chimerism, persistence of non-transformed escape tissues, or developmental changes in *GFP* detectability. Such factors are well recognized during prolonged citrus regeneration under selection, where non-transformed cells may persist alongside transformed tissues and contribute to escapes or chimeric shoots [11,17]. This underscores the value of visible markers: a signal that can be continuously and non-destructively monitored at each subculture may facilitate early identification of transformed sectors before escapes and chimeras become dominant. Visible pigmentation should therefore be viewed as a practical complement to antibiotic selection and fluorescence-based screening rather than as a replacement for molecular confirmation of regenerated plants.

### 3.5. Limitations

Several limitations should be considered when interpreting these findings. First, the transformation system was developed and validated in Carrizo citrange, a genotype with high regeneration competence; therefore, the performance of *ROSEA1* and *RUBY* should be evaluated in additional citrus genotypes and regeneration systems before broader application, since regeneration competence and marker expression in citrus are strongly genotype- and route-dependent [10, 11]. Second, differences in regeneration efficiency and callus biomass among treatments were secondary observations, and the experimental design was not intended to determine their underlying mechanisms. The higher regeneration observed in the reporter treatments cannot be definitively separated from enrichment of regeneration-competent tissues during continuous selection and therefore should be interpreted as evidence of the reporters’ compatibility with efficient shoot formation rather than evidence that they promote regeneration. Similarly, the contribution of pigment accumulation to increased callus biomass remains hypothetical because ROS levels and antioxidant status were not measured directly. Finally, the effects of constitutive reporter expression were evaluated only during *in vitro* regeneration, leaving potential impacts on subsequent plant growth and development unexamined, despite known developmental trade-offs associated with constitutive pigment expression in some species [25,50]. Future validation across additional citrus genotypes will be important to determine the broader applicability of the indirect organogenesis and visible-monitoring workflow developed here.

## 4. Materials and Methods

### 4.1. Plant material and seed sterilization

Carrizo citrange (*Citrus sinensis* × *Poncirus trifoliata*) seeds were obtained from Lyn Citrus Seeds (California, USA). Seeds were surface sterilized under aseptic conditions in a laminar airflow cabinet. Sterilization involved immersion in 70% (v/v) ethanol for 1 min, followed by 20% (v/v) commercial bleach solution (approximately 1% sodium hypochlorite) containing 2–3 drops of Tween-20 for 15 min with gentle agitation. Seeds were subsequently rinsed seven times with sterile distilled water to remove residual bleach and surfactant and were blotted dry on sterile filter paper prior to inoculation.

### 4.2. In vitro germination

Sterilized seeds were cultured on solid Murashige and Skoog (MS) basal medium [51] supplemented with 3% (w/v) sucrose and solidified with 0.7% (w/v) agar. The pH was adjusted to 5.8 prior to autoclaving at 121 °C for 15 min. Cultures were incubated in complete darkness at 25 °C for 6 weeks. Germinated seedlings were subsequently maintained under dark conditions for an additional 7 days to promote etiolation and epicotyl elongation. Etiolated seedlings served as the source of explants for transformation experiments.

### 4.3. Explant preparation

The epicotyl region (defined as the segment between the cotyledons and the first pair of true leaves) was excised from 7-week-old seedlings and sectioned into 5–7 mm segments using a sterile scalpel and forceps. To enhance *Agrobacterium* accessibility, both ends of each epicotyl segment were lightly wounded. Epicotyl segments were pre-cultured on callus induction medium (CIM0) without antibiotics for 5 days in complete darkness at 25 °C.

### 4.4. Vector construction

All constructs were based on the pOX135 binary vector [52], which carries a fused *eGFP–NPTII* gene under the control of the Cassava vein mosaic virus (CsVMV) promoter and terminated by the 35S terminator (35S-T), enabling both fluorescent detection and kanamycin selection, together with a spectinomycin resistance gene for bacterial selection. Coding sequences were introduced at the BsaI cloning sites under a 2×CaMV35S promoter containing an Omega enhancer (pGWB402, AB294426.1) and terminated by the heat-shock protein terminator (Hsp-T).

Two visible reporters were used. *ROSEA1* (GenBank AKB94073.1) encodes an R2R3-MYB transcription factor from *Antirrhinum majus* that activates anthocyanin biosynthesis through the endogenous MYB–bHLH–WD40 (MBW) complex [24]. *RUBY* is a synthetic cassette encoding three betalain biosynthetic enzymes linked by P2A self-cleaving peptides that together convert tyrosine to betalain [20]. The *ROSEA1* coding sequence was synthesized (Gene Universal Inc., Newark, DE, USA), and the *RUBY* cassette was amplified from plasmid 35S:*RUBY* (Addgene #160908), to generate *pOX135-ROSEA1* and *pOX135-RUBY*. The unmodified *pOX135* vector served as the control (CK).

Each construct was verified by PCR across the expression cassette (Fig. 1d) and confirmed by Sanger sequencing. Verified plasmids were introduced into *Agrobacterium* tumefaciens strain EHA105 by the freeze–thaw method, and transformed colonies were selected on LB medium supplemented with spectinomycin (100 mg L^-1^) and rifampicin (50 mg L^-1^), verified by colony PCR, and maintained as glycerol stocks at −80 °C.

### 4.5. Agrobacterium-mediated transformation

*Agrobacterium* tumefaciens strain EHA105 harboring the binary vectors was cultured in liquid yeast extract beef (YEB) medium supplemented with 100 mg L^-1^ spectinomycin and 25 mg L^-1^ rifampicin at 28 °C with shaking at 200 rpm overnight. A secondary culture was initiated by transferring 1 mL of the overnight culture into 10 mL of fresh LB broth containing the same antibiotics and grown until the OD600 reached 0.3–0.5 (mid-log phase). Bacterial cells were pelleted by centrifugation at 4,000 × g for 10 min and resuspended in infection buffer (½-strength MS salts supplemented with 10 mM MES, pH 5.8, 20 g L^-1^ sucrose, and 100 µM acetosyringone) to an OD_600_ of 0.2 for plant infection. Precultured epicotyl explants were immersed in the *Agrobacterium* suspension for 5 min with gentle agitation to ensure uniform contact. Following infection, explants were blotted dry on sterile filter paper and placed on co-cultivation medium (CIM supplemented with 100 µM acetosyringone and without antibiotic supplementation). Co-cultivation was carried out for 48 h in the dark at 25 °C.

### 4.6. Callus induction and selection

Following co-cultivation, explants were transferred to callus induction medium (CIM) to select for transformed tissue. CIM had the same composition as the pre-culture medium (CIM0) - full-strength MS salts and vitamins, 3% (w/v) sucrose, 1 mg L^-1^ 6-benzylaminopurine (BAP), 0.2 mg L^-1^ 1-naphthaleneacetic acid (NAA), and 0.5 mg L^-1^ 2,4-dichlorophenoxyacetic acid (2,4-D), pH 5.8, with the addition of 75 mg L^-1^ kanamycin to select transformed cells and 200 mg L^-1^ Timentin, to suppress residual *Agrobacterium*. Explants were maintained in darkness at 25 °C and subcultured onto fresh CIM every 2 weeks. Callus induction efficiency, callus fresh weight, and callus area were recorded at weeks 3 and 7 (Fig. 2). Callus induction efficiency was calculated as:

Callus induction efficiency (%) = Number of explants forming callus/Total number of explants cultured×100

Callus fresh weight was determined by excising callus from the explant and weighing immediately on an analytical balance and is reported as g per explant.

Callus area was measured from calibrated overhead images in ImageJ 1.46r (NIH, Bethesda, MD, USA) against an inframe scale reference and is reported as cm^2^ per explant.

### 4.7. Shoot regeneration and elongation

Actively growing callus was excised and transferred to shoot induction medium (SIM) containing MS salts and vitamins, 3% (w/v) sucrose, and 0.5 mg L^-1^ Thidiazuron (TDZ), 0.8% (w/v) agar, 2 mg L^-1^ silver nitrate, and 5% (v/v) coconut water, with the pH adjusted to 5.8, along with kanamycin (75 mg L^-1^) and Timentin (200 mg L^-1^). Developing shoots were sub-cultured onto fresh SIM every 2 weeks. Regenerated shoots (≥1 cm) were transferred to shoot elongation medium (SEM: MS salts and vitamins, 3% (w/v) sucrose, 0.5 mg L^-1^ BAP, 0.25 mg L^-1^ gibberellic acid (GA_3_), pH 5.8, with the same antibiotics) to promote elongation. Regeneration efficiency and shoots per explant were calculated as:

Regeneration efficiency (%) = Number of explants producing shoots / Total number of explants cultured×100 Shoots per explant = Total number of shoots / Total number of explants cultured

### 4.8. Rooting and Acclimatization

Elongated shoots were transferred to rooting medium (½-strength MS salts, 2% (w/v) sucrose, 0.5 mg L^-1^ indole-3- butyric acid (IBA), pH 5.8) under a 16 h photoperiod at 25 °C. The medium was supplemented with kanamycin (75 mg L^-1^) and Timentin (200 mg L^-1^). Successfully rooted plantlets were gradually acclimatized by transferring them to sterile substrate under high humidity and subsequently maintained under controlled greenhouse conditions. Rooting efficiency was calculated as:

Rooting efficiency (%) = Number of shoots forming roots / Total number of shoots transferred to rooting medium × 100

### 4.9. Assessment of transformation efficiency

Transformation efficiency was assessed using both GFP fluorescence and visible pigmentation. GFP fluorescence was detected using a fluorescence stereomicroscope (Leica M205 FA) with a GFP filter set; pigmentation was scored macroscopically under ambient light. Scoring was performed at the callus stage at weeks 3 and 7, and at the shoot stage at weeks 13 and 17. Transformation efficiency at each stage was calculated per criterion as:

Transformation efficiency callus (%) = Number of explants positive for *GFP* (or pigment) / Total number of explants cultured × 100

Transformation efficiency shoot (%) = Number of shoots positive for *GFP* (or pigment) / Total number of regenerated shoots × 100

### 4.10. Anthocyanin and betalain quantification

Anthocyanin content was determined spectrophotometrically following established protocols [53]. Approximately 100 mg of callus tissue was homogenized in 1 mL of acidified methanol (1% HCl) and incubated overnight at 4 °C in the dark. Samples were centrifuged at 12,000 × g for 10 min, and the absorbance of the supernatant was measured at 530 nm and 657 nm using a UV-visible spectrophotometer (NanoDrop 2000).

Anthocyanin content was calculated using the formula: A_530_ − 0.25 × A_657_, and expressed as mg g^-1^ fresh weight (FW).

Total betalain content was quantified as described by [54]. Callus tissue (100 mg) was homogenized in distilled water and centrifuged at 12,000 × g for 10 min. Absorbance of the supernatant was measured at 538 nm (betacyanins) and 480 nm (betaxanthins). Total betalain content was calculated and expressed as mg g^-1^ FW.

Betalain content was calculated using the formula: Betalain content (mg/g FW) = (A × DF × MW × V) / (ε × L × W), where A = absorbance, DF = dilution factor, MW = molecular weight (550 g/mol for betacyanins; 308 g/mol for betaxanthins), V = extract volume (mL), ε = molar extinction coefficient (60,000 L/mol·cm for betacyanins; 48,000 L/mol·cm for betaxanthins), L = path length (1 cm), and W = fresh weight (g) [55].

### 4.11. Molecular confirmation of transformants

Putative transformation events growing under kanamycin selection media were further validated via PCR. For this, Genomic DNA was isolated from 100 mg of leaf tissue using a modified cetyltrimethylammonium bromide (CTAB) method [56]. Primers targeting the genes *ROSEA1* and *RUBY* (supplementary Table S1) were used to screen the putative transformants, using the GoTaq® Green Master Mix (Promega) on the BioRad thermocycler conditions listed in supplementary Table S1. Resulting amplicons were visualized in 1% agarose gel stained with SYBR Safe DNA Gel Stain on the Gel-Doc imager (Thermo Fisher Scientific, USA). Wild-type (non-transformed) Carrizo served as the negative control, and plasmid DNA served as the positive control.

### 4.12. Experimental design and statistical analysis

Each transformation experiment was performed with three treatments - the control vector (CK), *pOX135-ROSEA1* (*ROS1*), and *pOX135-RUBY* - and repeated at least three times. A biological replicate was defined as one culture plate, with 8–10 replicates and a total of 50–60 explants per treatment. Unless otherwise indicated, experimental week numbers were counted from the transfer of co-cultivated explants to selective CIM, which was designated as week 0. Callus induction efficiency, callus fresh weight, and callus area were recorded at weeks 3 and 7; regeneration efficiency and shoots per explant at weeks 5, 7, and 9; and callus and shoot transformation efficiency, scored by GFP fluorescence and by visible pigmentation, at weeks 3 and 7 and weeks 13 and 17, respectively. Anthocyanin content was quantified in CK and *ROSEA1* tissues, and betalain content in CK and *RUBY* tissues, at weeks 3 and 7.

Data are presented as mean ± standard deviation (s.d.). Statistical significance was assessed using two-tailed Student’s t-tests (CK versus *ROSEA1*; CK versus *RUBY*). A p-value < 0.05 was considered statistically significant. All analyses were performed in R (version 4.3.2), and figures were generated using ggplot2.

## 5. Conclusions

In this study, we established an *Agrobacterium*-mediated indirect organogenesis-based transformation system for Carrizo citrange that supported callus induction, shoot regeneration, rooting, and recovery of PCR-confirmed transgenic plantlets within approximately 20 weeks. Integration of *ROSEA1* and *RUBY* visible reporters enabled transformed tissues to be monitored non-destructively from callus proliferation through plant recovery, with *GFP* providing an independent fluorescence-based reference. Both reporters performed effectively during the callus stage, while *RUBY* maintained more consistent visual detectability during shoot regeneration than *ROSEA1*. Importantly, neither reporter prevented efficient regeneration or recovery of transgenic plants. Overall, this work provides a practical indirect organogenesis-based platform for citrus transformation and demonstrates the value of visible pigmentation as an equipment-free complement to antibiotic selection and fluorescence-based screening during prolonged regeneration. The system provides a useful foundation for future citrus functional genomics and genome-editing applications.

## Supplementary Materials

The following supporting information are provided in the accompanying supplementary file.

Supplemental Figure S1: Representative citrus explants during *Agrobacterium*-mediated transformation at week 3. Additional biological replicates of citrus explants under control (CK), *ROSEA1* (*ROS1*), and *RUBY* treatments at week 3. Five representative plates per treatment are shown. Scale bars, 1 cm.

Supplemental Figure S2: Representative citrus explants during *Agrobacterium*-mediated transformation at week 5. Additional biological replicates of citrus explants under control (CK), *ROSEA1* (*ROS1*), and *RUBY* treatments at week 5. Five representative plates per treatment are shown. Scale bars, 1 cm.

Supplemental Figure S3: Representative citrus explants during *Agrobacterium*-mediated transformation at week 7. Additional biological replicates of citrus explants under control (CK), *ROSEA1* (*ROS1*), and *RUBY* treatments at week 7. Five representative plates per treatment are shown. Scale bars, 1 cm.

Supplemental Figure S4: Representative citrus explants during *Agrobacterium*-mediated transformation at week 9. Additional biological replicates of citrus explants under control (CK), *ROSEA1* (*ROS1*), and *RUBY* treatments at week 9. Five representative plates per treatment are shown. Scale bars, 1 cm.

Supplemental Figure S5: Representative citrus explants during *Agrobacterium*-mediated transformation at week 15. Additional biological replicates of citrus explants under control (CK), *ROSEA1* (*ROS1*), and *RUBY* treatments at week 15. Four representative plates per treatment are shown. Scale bars, 1 cm.

Supplemental Figure S6: Representative fully developed wild-type citrus plantlets at week 20. Whole-plant overview (left), close-up leaf images (middle), and GFP fluorescence (right) imaging of leaves are shown. Scale bars, 1 cm.

Table S1: The list of primers used in this study.

## Author Contributions

T.J. and S.E.T. conceived and designed the study. S.E.T. performed the experiments. T.J. and S.E.T. conducted data analysis and visualization. S.E.T. and T.J. wrote the manuscript with input from all authors. H.H., C.D.M., T.J., and A.K. revised the manuscript. All authors have read and agreed to the published version of the manuscript.

## Funding

This work was supported by USDA-NIFA (grant number 2019-67013-29236) and the USDA HATCH program (grant number FLA-MFC-006387) to H.H.

## Data Availability Statement

All data supporting the findings of this study are available from the corresponding author upon reasonable request.

## Acknowledgments

We thank Jaideep Chandranshu Cherukula for plant and facility management and Keila Emily Rodriguez for the transgenic lines previously generated for this project.

## Conflicts of Interest

The authors declare no conflicts of interest.

## Abbreviations

The following abbreviations are used in this manuscript:

*BAP*: 6-Benzylaminopurine
*CCM*: Co-cultivation medium
*CIM*: Callus induction medium
*CLas*: Candidatus Liberibacter asiaticus
*eGFP*: Enhanced green fluorescent protein
*HLB*: Huanglongbing
*IBA*: Indole-3-butyric acid
*NAA*: 1-Naphthaleneacetic acid
*NPTII*: Neomycin phosphotransferase II
*RIM*: Rooting medium
*SEM*: Shoot elongation medium
*SIM*: Shoot induction medium
*TDZ*: Thidiazuron

## Notes

### Competing Interest Statement

The authors have declared no competing interest.

