## Supplementary for "An indirect organogenesis-based citrus transformation system monitored using visible reporter markers"

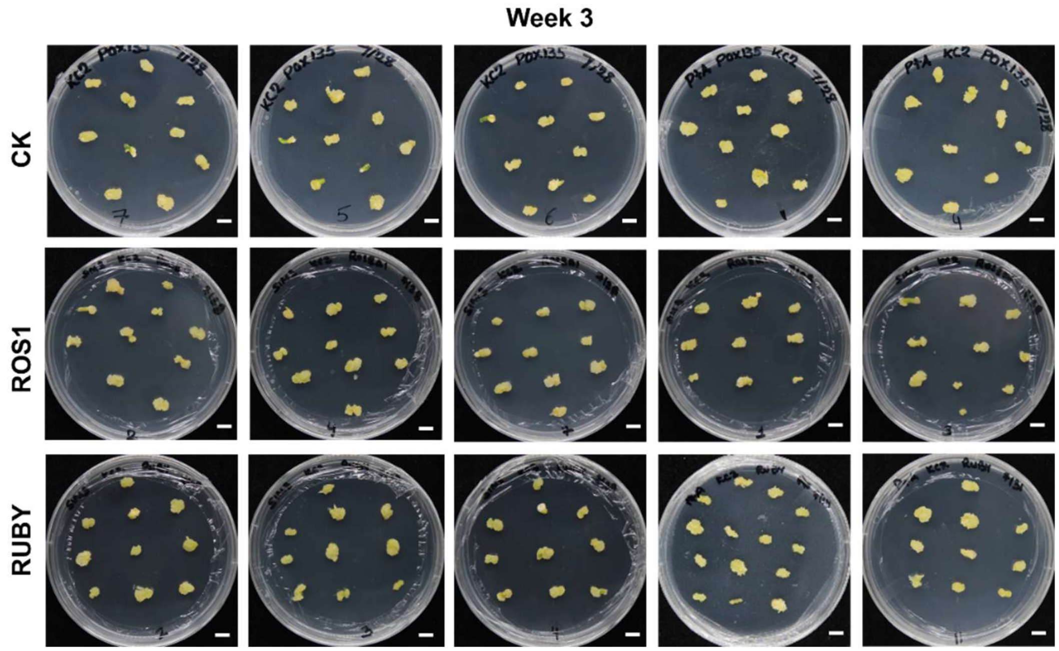


**Supplemental Fig. S1 | Representative citrus explants during *Agrobacterium*-mediated transformation at week 3.** Additional biological replicates of citrus explants under control (CK), *ROSEA1* (*ROS1*), and *RUBY* treatments at week 3. Five representative plates per treatment are shown. Scale bars, 1 cm.

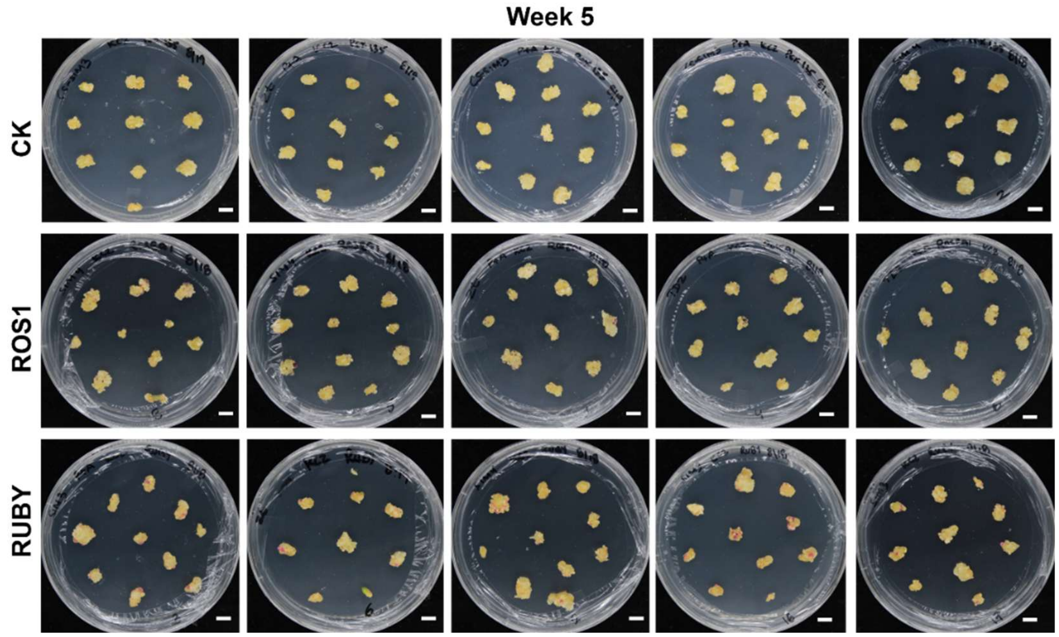


**Supplemental Fig. S2 | Representative citrus explants during *Agrobacterium*-mediated transformation at week 5.** Additional biological replicates of citrus explants under control (CK), *ROSEA1* (*ROS1*), and *RUBY* treatments at week 5. Five representative plates per treatment are shown. Scale bars, 1 cm.

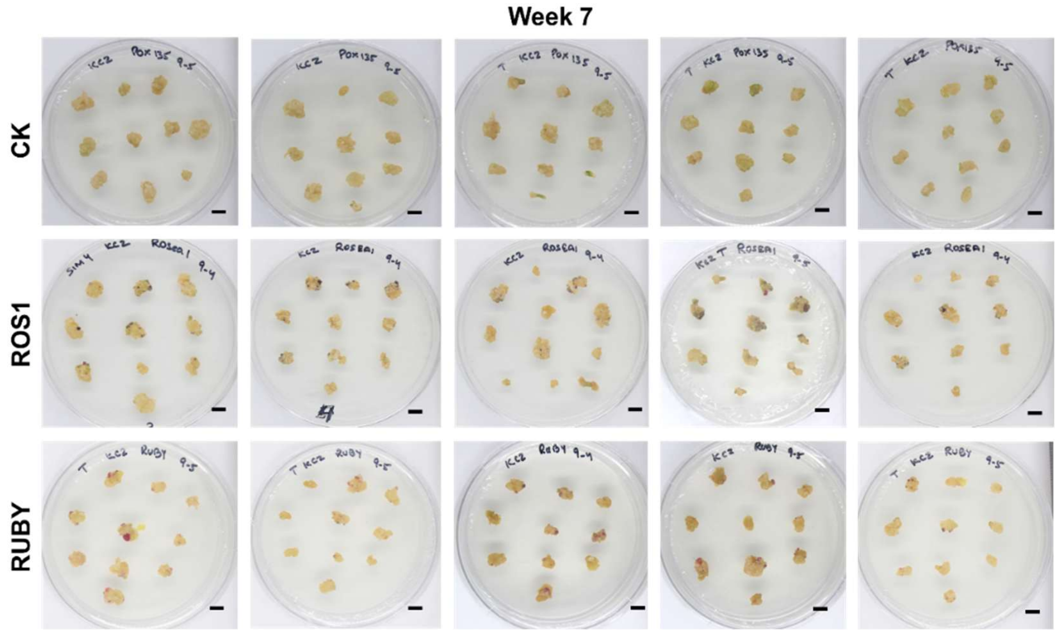


**Supplemental Fig. S3 | Representative citrus explants during *Agrobacterium*-mediated transformation at week 7.** Additional biological replicates of citrus explants under control (CK), *ROSEA1* (*ROS1*), and *RUBY* treatments at week 7. Five representative plates per treatment are shown. Scale bars, 1 cm.

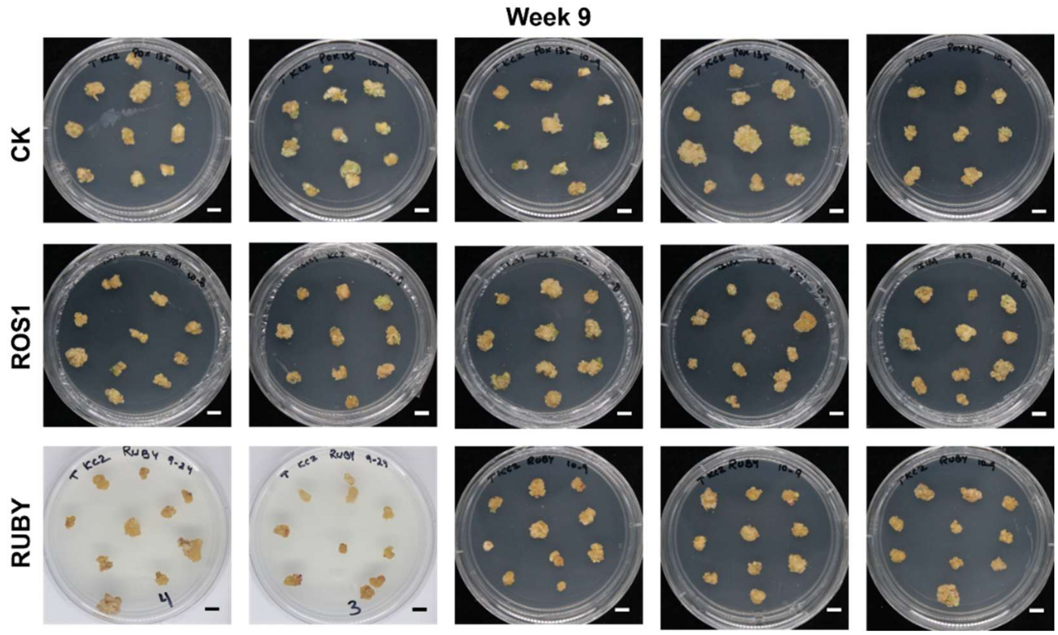


**Supplemental Fig. S4 | Representative citrus explants during *Agrobacterium*-mediated transformation at week 9.** Additional biological replicates of citrus explants under control (CK),

*ROSEA1* (*ROS1*), and *RUBY* treatments at week 9. Five representative plates per treatment are shown. Scale bars, 1 cm.

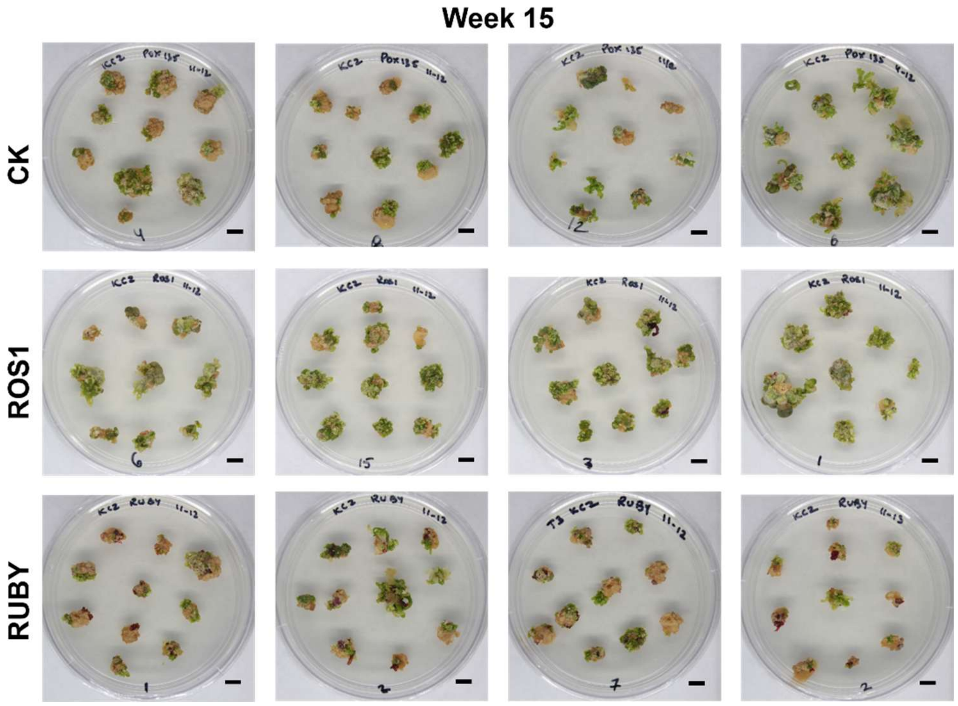


**Supplemental Fig. S5 | Representative citrus explants during *Agrobacterium*-mediated transformation at week 15.** Additional biological replicates of citrus explants under control (CK), *ROSEA1* (*ROS1*), and *RUBY* treatments at week 15. Four representative plates per treatment are shown. Scale bars, 1 cm.

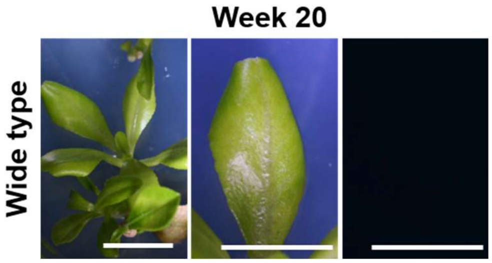


**Supplemental Fig. S6 | Representative fully developed wild-type citrus plantlets at week 20.** Whole-plant overview (left), close-up leaf images (middle), and GFP fluorescence (right) imaging of leaves are shown. Scale bars, 1 cm.

| Description | Sequence | Amplicon size (bp) | Annotation |
|---|---|---|---|
| *RUBY-F* | CACGAACTCCAGCAGGACCATG | 630 | Transgenic plant genotyping PCR |
| *RUBY-R* | CTGGTCGAGCTGGACGGCGACG | | |
| *AmROSEA1*-F | ATGGAAAAGAATTGTCGTGG | 450 | |
| *AmROSEA1*-R | TTAATTTCCAATTTGTTGGGC | | |
| *Insert-dec-F* | atgaattCCAACATGGTGGAGCAC | CK: 1.1 kb; ROSEA1: 1.7 kb; RUBY: 3.2 kb | Plasmid diagnostic PCR |
| *Insert-dec-R* | TGCTGCAGAAGAGATCCAACA | | |

**Supplemental Table 1. The list of primers used in this study**
